# Optogenetic control of YAP can enhance the rate of wound healing

**DOI:** 10.1101/2022.11.04.515183

**Authors:** Pearlyn Jia Ying Toh, Marius Sudol, Timothy Edward Saunders

**Affiliations:** Mechanobiology Institute, National University of Singapore, Singapore; Icahn School of Medicine at Mount Sinai, New York City, USA; Institute of Molecular and Cell Biology, A*STAR, Singapore; Warwick Medical School, University of Warwick, United Kingdom

**Keywords:** Hippo-YAP, optogenetics, wound healing

## Abstract

**Background:** Tissues need to regenerate to restore function after injury. Yet, this regenerative capacity varies significantly between organs and between species. For example, in the heart, some species retain full regenerative capacity throughout their lifespan but human cardiac cells display limited ability to repair injury. After a myocardial infarction, the function of cardiomyocytes is impaired and reduces the ability of the heart to pump, causing heart failure. Therefore, there is a need to restore the function of an injured heart post myocardial infarction. We investigate in cell culture the role of the YAP, a transcriptional co-regulator with a pivotal role in growth, in driving repair after injury.

**Methods:** We express optogenetic YAP (optoYAP) in three different cell lines. We characterised the behaviour and function of optoYAP using fluorescence imaging and quantitative real-time PCR of downstream YAP target genes. Mutant constructs were generated using site-directed mutagenesis. Nuclear localised optoYAP was functionally tested using wound healing assay and anchorage-independent colony formation assay.

**Results:** Utilising optoYAP, which enables precise control of pathway activation, we show that YAP induces the expression of downstream genes involved in proliferation and migration. optoYAP can increase the speed of wound healing in H9c2 cardiomyoblasts. Interestingly, this is not driven by an increase in proliferation, but by collective cell migration. We subsequently dissect specific phosphorylation sites in YAP to identify the molecular driver of accelerated healing.

**Conclusions:** This study shows that optogenetic YAP is functional in H9c2 cardiomyoblasts and its controlled activation can potentially enhance wound healing in a range of conditions.

## Introduction

During myocardial infarction, blood flow to the heart is severely limited or completely cut off by plaque or cholesterol blockages. Without oxygen and nutrients, heart muscles surrounding the region of the blocked vessel begin to necrose. Acute myocardial injury can kill 25% of cardiomyocytes, rendering them non-functional (1). Pharmacological management after a heart attack only serves to prevent another episode from occurring and there are no known strategies to restore the full capacity of the heart. This is in part due to the human heart having a very limited ability to regenerate post injury (2) and the lack of regenerative capability is in contrast to other organs, such as the liver or gut (3). Further, other organisms show a greater capacity to regenerate injured heart tissue post-embryonic development: young mice (<7 days) can repair heart wounding; and in zebrafish the potential to regenerate heart tissue remains throughout the lifespan (4,5). Therefore, there is a demand to find strategies to induce cardiac regeneration in humans. Though new cardiomyocytes can be synthesised *ex vivo* from embryonic stem cells and organise to form contractile tissues, injected cells have an embryonic lineage and can generate arrythmias (6,7). Recipients also need to be immunosuppressed for transfusion, which itself is a complex procedure. Alternative strategies to induce human heart tissue repair and regeneration are still heavily sought after.

A strategy to circumvent the above complexity is to stimulate the endogenous proliferation potential of cells in repair. The key idea is to effectively mimic developmental processes, reviving pathways that are switched off postnatally. This approach has seen positive results in transgenic mice models, where activated YAP promotes cardiac regeneration by inducing reparative genetic programs to re-enter the cell cycle and activate proliferative genes in cardiomyocytes (8–10). The Hippo-YAP pathway is highly implicated in development and regenerative processes, due to its role in driving cell proliferation, migration and mechanosensing (11,12). The pathway is constructed principally from a kinase cascade that culminates at the transcriptional co-activator YAP and its paralogue TAZ. Phosphorylated YAP/TAZ triggers its cytoplasmic sequestration (13,14) or degradation (15), while unphosphorylated YAP/TAZ can translocate into the nucleus and bind to transcription factors TEAD1-4 (16–18). This stimulates the expression of a range of genes that typically lead to pro-survival outcomes in the cell (19–21). The nucleocytoplasmic distribution of YAP/TAZ is a key determinant of its activity. Therefore, manipulating the YAP/TAZ subcellular localisation can provide a tool to control the activation of the Hippo-YAP pathway.

Previous studies have shown that the MAPK/ERK pathway is a key regulator of organ regeneration (22,23), and crosstalks with the Hippo-YAP pathway in cardiomyocytes reactivates cell proliferation (24,25). ERBB2 (a member of the epidermal growth factor receptor (EGFR) family) signalling induces a phosphorylation mark on two non-canonical serine residues on YAP by ERK (25). The activating phosphorylation marks on S251 and S333 oppose the conventional deactivating phosphorylation caused by LATS on S127. The phosphorylation mark at S127 generates a 14-3-3 binding site, causing the cytoplasmic sequestration of phosphorylated YAP, accelerating the nuclear export of YAP and inhibiting its co-transcriptional function inside the nucleus (26,27). Mutation of serine 127 to alanine has been reported to increase the activity of YAP by preventing LATS phosphorylation (28–30). Besides the phosphorylation site at S127, another protein domain that regulates the localisation of YAP is the C-terminal PDZ-binding motif (31), a short motif of only five amino acid residues, -FLTWL at the very carboxyl terminal end. The PDZ-binding motif allows YAP to bind to proteins containing the PDZ domain such as ZO-2, and a deletion of this motif impairs the nuclear shuttling of YAP through ZO-2 (31). However, similar to the S127A mutant, truncation of the PDZ-binding motif does not impair the induction of TEAD-mediated transcriptional activity, only affecting its subcellular localisation (30,32). The YAP paralogue TAZ also contains a C-terminal PDZ-binding motif, which has been shown to be required for its nuclear localisation (14). Therefore, the PDZ-binding motif appears to be a conserved sequence that is integral to the function of YAP/TAZ as transcriptional co-regulators. However, how these phosphorylation marks and domains alter the dynamics of YAP signalling – and subsequently its efficacy in regeneration – remain unknown.

During regeneration, there are two main mechanisms for repair: (i) migration of newly proliferated cells into the injured area; and (ii) cell proliferation within the regenerating zone. YAP is involved in wound healing by limiting cytoskeletal and focal adhesion maturation to promote cell motility (33). YAP increases migration and invasiveness in a number of cell lines, often stimulating epithelial-to-mesenchymal transition (EMT) (34). On top of enhancing cell cycle progression, YAP also promotes cytoskeletal remodelling, responding to the mechanical change during scar tissue formation post myocardial infarction (35). YAP has also been shown to be activated in cardiac fibroblasts after ischemic injury (36).

While YAP activity has been shown to regenerate the mice heart after injury, caution must be taken as over-proliferation can cause occlusive vascular diseases (37). Therefore, having control over the spatiotemporal localisation of YAP is important. Optogenetics provides such a method (38,39), whereby light activation can control protein behaviour, and we have recently applied this to YAP, generating an optogenetic YAP construct termed optoYAP (40). Can optogenetic YAP be used to control regeneration after wounding or injury? optoYAP is potentially a powerful tool to circumvent regenerative limitations whilst retaining specific spatiotemporal control over its activity. Building upon our previously reported work (40), we utilise a range of cell lines, including H9c2 rat cardiomyoblasts, as *in vitro* models to study how activation of YAP can promote repair post-injury (41,42)

Here, we report that activated optoYAP accelerates wound healing in two different cell lines, driving cell migration that is essential during regeneration. In particular, we show that H9c2 cardiomyoblasts are responsive to optoYAP activation, inducing nuclear accumulation after light activation. Nuclear optoYAP increases expression of downstream proliferative YAP target genes *CTGF* and *CYR61*, as well as *TGF-β*, a transforming growth factor involved in stem cell regulation and differentiation. This suggests that optoYAP can potentially be applicable in regenerative studies of tissues – including cardiac tissues – by promoting the renewal and replication of cells post-injury.

Crosstalk between the Hippo-YAP and ERBB2 signalling pathways during cardiac regeneration generates two activating phosphorylations by ERK at S251 and S333 on YAP (25). To further understand the potential mechanism for YAP function in regeneration, we generated phosphomimetic mutants at both serine residues and observed that they had an elevated amount of nuclear localisation in the dark as compared to wildtype optoYAP. However, the mutants could still respond to light activation. Building upon the changes in basal distribution of mutant optoYAP, we mutated the canonical LATS phosphorylation serine residue 127 to alanine, preventing its inactivating phosphorylation effect on YAP. We observed that optoYAP S127A was strongly nuclear localised before light activation and the nuclear signal was amplified after light activation. By removing the C-terminal PDZ-binding motif, another regulatory domain of YAP, we arrested the intrinsic nuclear localisation of YAP by preventing it from binding to nuclear importins. However, optogenetic activation was still able to promote nuclear localisation of this mutant through the nuclear localisation sequence (NLS) present on the optogenetic backbone. Overall, our results reveal that optogenetic approaches provide a strategy for specific control over YAP nuclear localisation (and hence activity) in both wildtype YAP and its mutants, thereby making it a potential tool for driving tissue regeneration.

## Materials and Methods

### Plasmid construction

Optogenetic plasmid containing YAP (optoYAP) was previously described by Toh et al., 2022 (40). Mutant YAP containing S127A or ΔC modifications were PCR amplified from pBABE (hygro) hYAP1-1δ S127A and pBABE (hygro) hYAP1-1δ ΔC plasmids respectively, both of which were from the library of YAP complementary DNA (cDNA) constructs of the Sudol laboratory. These constructs are available from ‘Addgene’ vector resource. Phusion High Fidelity DNA polymerase (ThermoScientific) was used for PCR amplification with primer sequences for each amplification listed in Table 1. PCR products and optoYAP plasmid were cleaved with appropriate FastDigest restriction enzymes (ThermoScientific) for cloning.

**Table 1.**
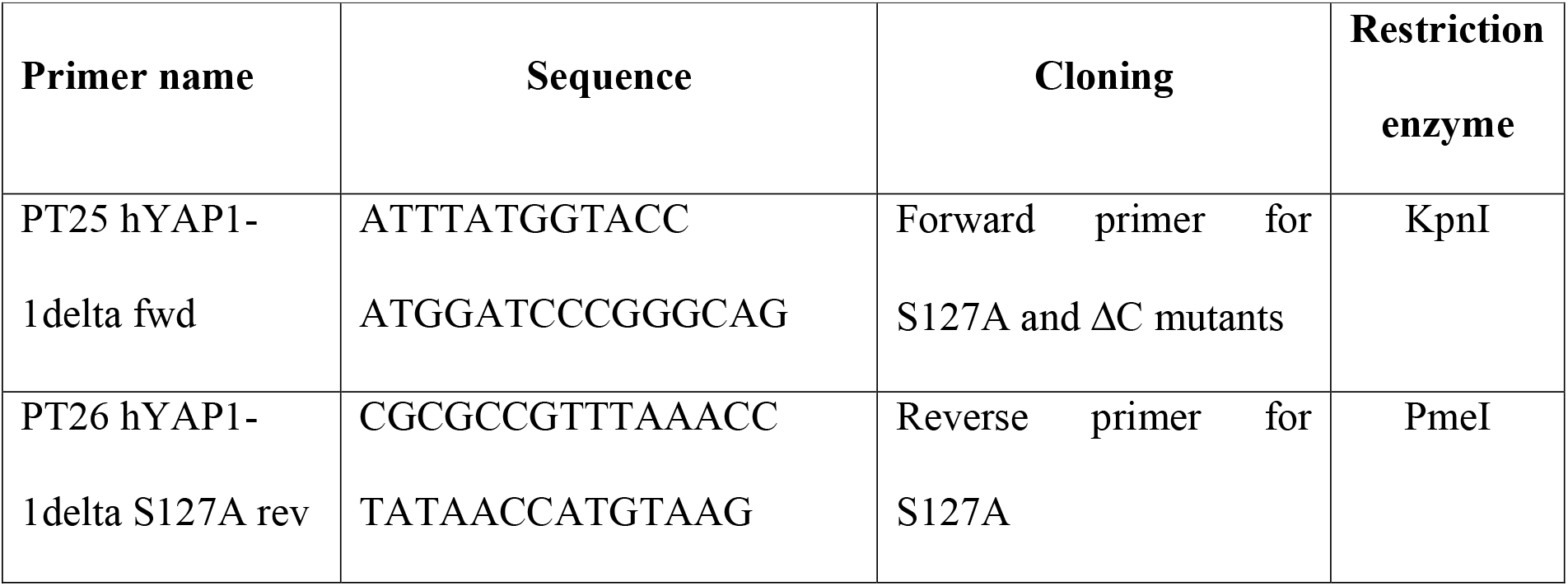
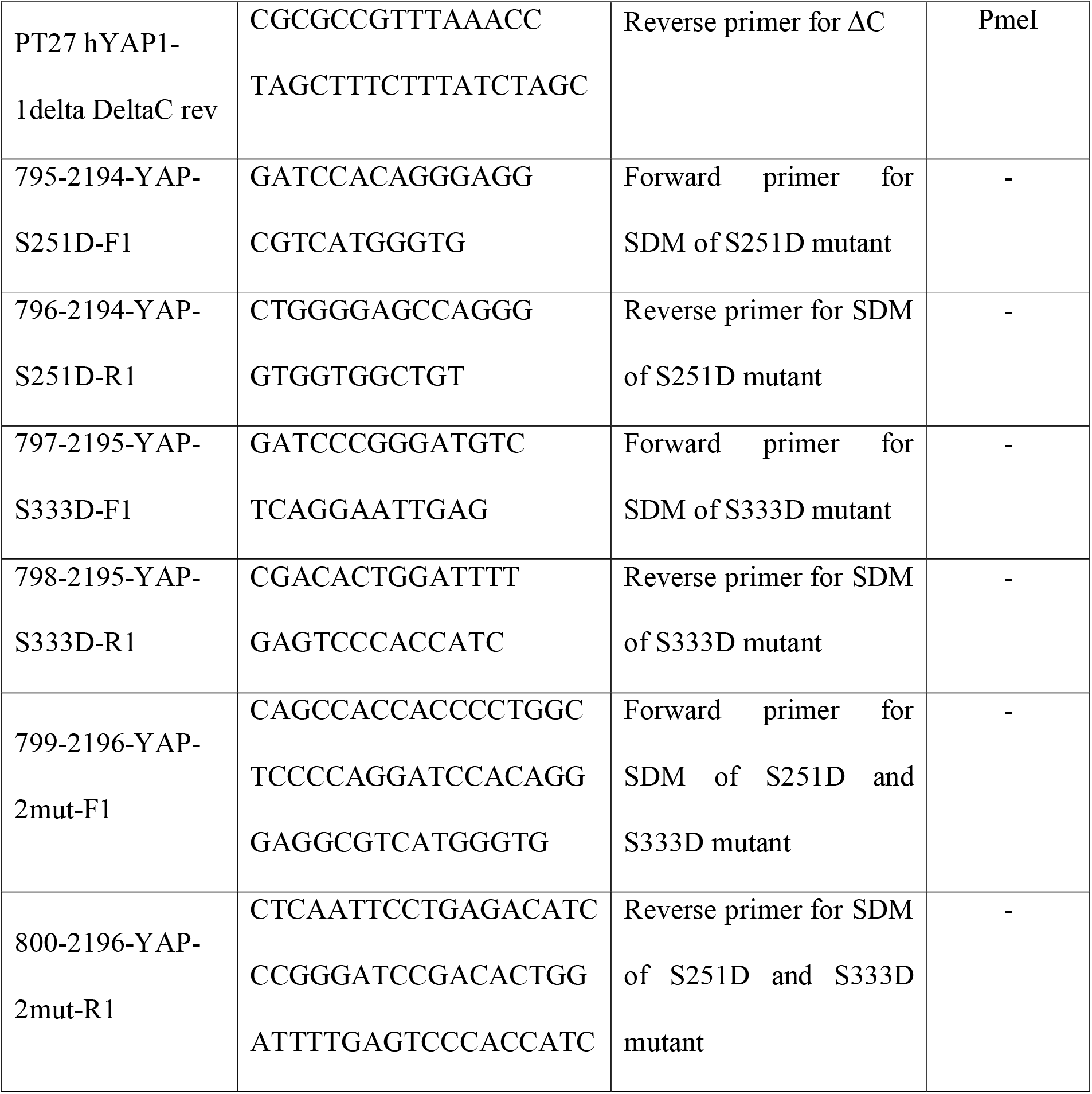
List of primers for plasmid cloning.

Point mutations in hYAP1-1δ were generated with Q5 Site-Directed Mutagenesis Kit (NEB) according to the manufacturer’s instructions to synthesise phosphomimic mutants by substituting serine with aspartic acid at two residues, S251 and S333. Positive clones were verified by restriction enzyme digestion and sequencing. Primers used for site-directed mutagenesis (SDM) are listed in Table 1.

### Mammalian cell culture and transfection

HEK293T (ATCC CRL-3216) human embryonic kidney cells were grown in Dulbecco’s modified Eagle’s medium (DMEM, ThermoScientific) supplemented with 10% (v/v) heat inactivated fetal bovine serum (FBS, Gibco) and 1% (v/v) penicillin/streptomycin (Gibco). MKN28 (CVCL_1416) human gastric adenocarcinoma cells were grown in RPMI (ThermoScientific) supplemented with 10% (v/v) FBS and 1% (v/v) penicillin/streptomycin. H9c2 (ATCC CRL-1446) rat cardiomyocytes (a gift from Dr Manvendra K. Singh’s lab) were grown in DMEM supplemented with 15% (v/v) FBS, 1% (v/v) penicillin/streptomycin and 1% non-essential amino acids (NEAA, Gibco). All cell cultures were kept in a 37ºC incubator with 5% CO_2_ and passaged every 2-3 days. Cells were routinely checked for mycoplasma contamination and short tandem repeat (STR) profiled (1^st^ Base).

HEK293T cells were transfected with plasmid DNA using Lipofectamine 2000 (ThermoScientific) using 6 μL of Lipofectamine 2000 reagent in 150 μL opti-MEM (ThermoScientific) and 3.5 μg of plasmid DNA in 175 μL opti-MEM according to the manufacturer’s instructions. H9c2 cells were transfected with Lipofectamine 2000 as per HEK293T cells, but with 5 μg of plasmid DNA. MKN28 cells were transfected using K2 transfection system (Biontex) according to the manufacturer’s instructions.

### Fluorescence microscopy

Live imaging was performed on Perkin Elmer spinning disk with a LUCPlanFLN 40x/0.6 numerical aperture (NA) air objective or Nikon Ti-E with Yokogawa W1 spinning disk with a PLAN APO VC 60x/1.20 NA water immersion objective. Cells were visualised under the microscope 24 h post transfection in a 37ºC heated and humidified chamber.

Long term imaging of wound healing experiments was performed on widefield Nikon Biostation IMQ with 10x air objective or Olympus IX81 with a UPLFLN 10x/0.3 NA air objective. The samples were tracked every 30 min to measure the rate of wound closure until the gap was completely closed.

### Image analysis

The images were analysed with Fiji ImageJ software. The nuclear to cytoplasmic ratio was measured by drawing a region of interest (ROI) around the cell as well as its nucleus. A macro plugin written by Richard De Mets was used to mask the nucleus from the cytoplasm to calculate the mean intensity in the nucleus and cytoplasm based on the ROI drawn. The ROI was drawn for each cell analysed at every time frame. The wound healing analysis was performed using a macro written by Ong Hui Ting, which measures the changes in gap area of the wound over time. Charts were made with Prism 8 software. All figures were assembled with Adobe Illustrator.

### Light activation

Pulsed light activation followed a protocol of 1 s pulse of 0.15 mW 488 nm laser light followed by a 30 s dark phase. This pulsation is continued for 20 min, which constitutes as ‘light activation’. During recovery, the entire set up is kept in the dark for a further 20 min. Throughout the entire 40 min of light activation and recovery, imaging was done with a 561 nm laser line to image the mCherry fluorescent protein. For samples kept under dark conditions, aluminium foil was used to wrap the sample to prevent light contamination.

### Quantitative real-time PCR (RT-qPCR)

Transfected cells were subjected to 48 h of pulsatile light activation and total RNA was extracted from cells using RNeasy Plus Mini Kit (QIAGEN) and QIAshredder (QIAGEN) according to the manufacturer’s instructions. RNA was eluted with 40 μL nuclease-free water (NFW) and quantitated using NanoDrop 2000 spectrophotometer. cDNA was synthesised using SuperScript IV Reverse Transcriptase (ThermoScientific) using oligo d(T) primers according to the manufacturer’s instructions.

Real-time PCR detection were done using SYBR Green PCR Master Mix for 40 cycles in a Bio-Rad CFX96 thermal cycler. The threshold cycle (Ct) value for each gene was normalised to the Ct value of a housekeeping gene, *EIF1B*. The relative fold changes were calculated using ΔΔCt method. The primer sequences for target genes are listed in Table 2.

**Table 2.**
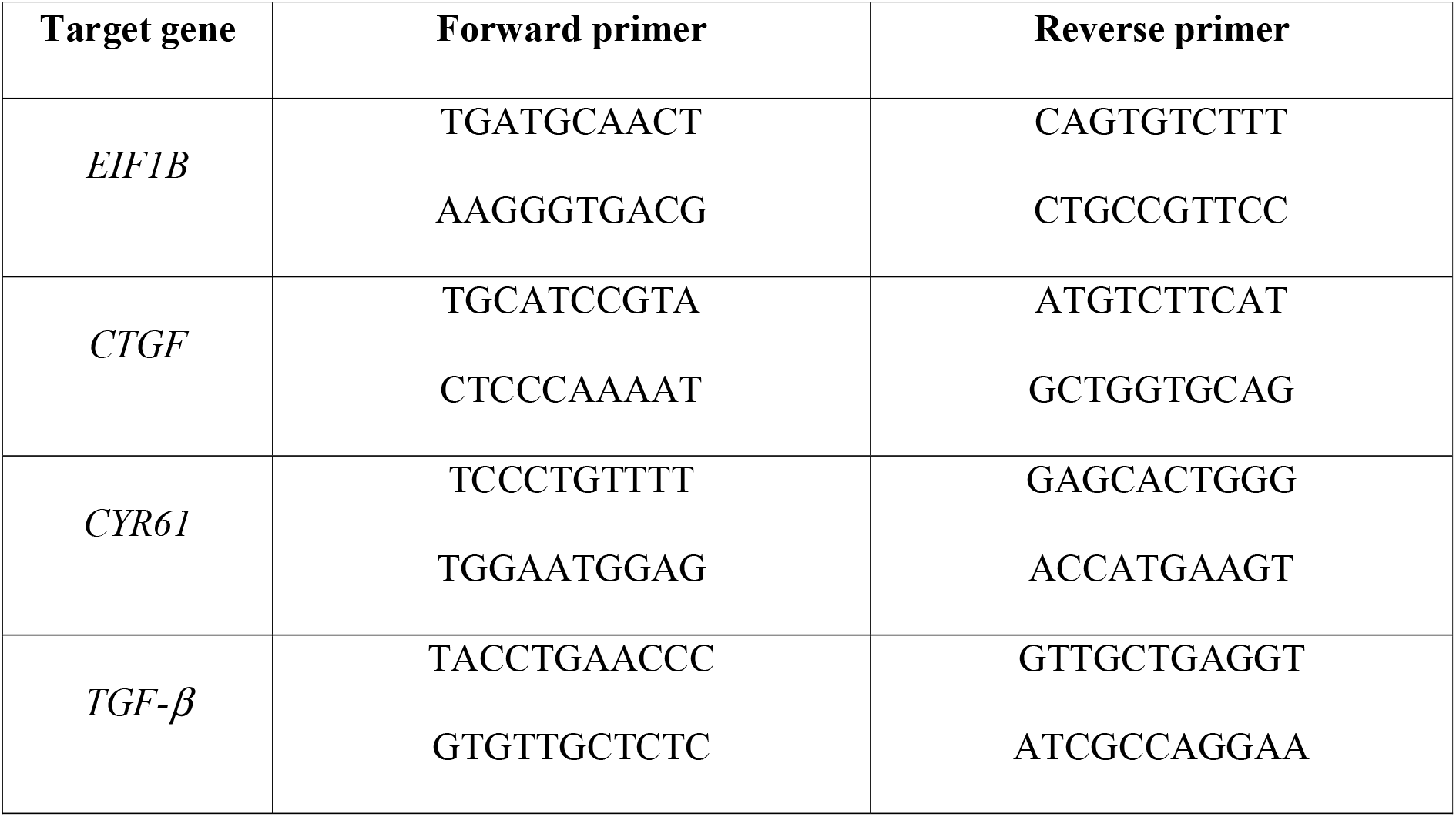
Primers used for RT-qPCR.

### Wound healing assay

Cells were grown in their respective serum free media for 24 h before 25000 cells per chamber were plated into Culture-Insert 2 well in μ-Dish 35 mm, high (ibidi). Cells were kept in serum free media throughout the experiment. 24 h after seeding, or when cells reach confluency in the chambers, the culture insert was removed with clean forceps under sterile conditions and the dish was filled with 2 mL of serum free media for imaging.

### Anchorage-independent assay

Anchorage-independent assay was performed using CytoSelect 96-Well Cell Transformation Assay (Cell Biolabs) according to the manufacturer’s instructions. Cells were grown in T25 flasks to 80% confluency, trypsinised and counted with a hemocytometer to obtain the concentration of live cells. Each well was seeded with 5000 cells and the final cell culture agar layer solution was 0.4% for soft conditions and 1.2% for stiff conditions, corresponding to a Young’s modulus of <2 kPa (43) and 10 kPa (44) respectively. Fresh medium was changed every 2 days during incubation and 7 days after seeding, cell colony formation was examined with EVOS Cell Imaging System (ThermoScientific) under 10x magnification. The total amount of DNA present in each well was measured with a DNA binding fluorescent dye using the CyQuant NF Cell Proliferation Assay Kit (ThermoScientific) according to the manufacturer’s instructions.

### Statistical analysis

Biological replicates refer to independent experimental replicates done on different days with fresh biological samples and independent transfections and measurements. Technical replicates refer to repeated measurements of the same sample that represent independent measures of the random noise associated with protocols or equipment (45). For comparisons between the same sample in different light and dark conditions, a paired t-test was used. For comparisons between two distinct samples, an unpaired t-test was used. For small sample sizes, estimation statistics was used to show the magnitude of the testing condition (46).

### Curve fitting

Curve fitting for the import and export rates was performed in MATLAB using the curve fitting function *fit*. Time constant (τ) represents the time scale over which the signal changes before or after light activation. Nuclear import rate (τ_a_) was fitted with 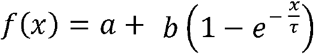 and export rate (τ_r_) was fitted with 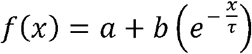, where *a* is the basal level of signal intensity; and *b* is the multiplicative factor representing the change in signal due to light (de-)activation.

## Results

### optoYAP activity in H9c2 cells

Using a previously reported optoYAP for optogenetic control of YAP nuclear import and export (40), we transfected H9c2 cells with optoYAP and characterised the construct in the rat cardiomyoblasts. We subjected transfected cells to pulsatile light activation with 488 nm laser power at 0.15 mW for 1 s at a 30 s interval, for a total of 20 min. We observed nuclear localisation of the optogenetic construct after the 20 min activation window (Fig. 1A). After leaving the cells in the dark for a further 20 min, optoYAP translocated out of the nucleus and returned to the basal state before light activation (Fig. 1A). The nuclear/cytoplasmic ratio of optoYAP almost doubled after the 20 min activation (Fig. 1B). Throughout the activation and recovery cycle, we tracked and imaged the cells over time to obtain the nuclear import and export dynamics (Fig. 1B). The activation (τ_a_) and recovery (τ_r_) time constants were calculated to be 4.6 ± 0.6 min and 13.5 ± 1.6 min respectively (Methods). These values are comparable with those obtained by using the HEK293T cell line described previously (40).

**Figure 1.**
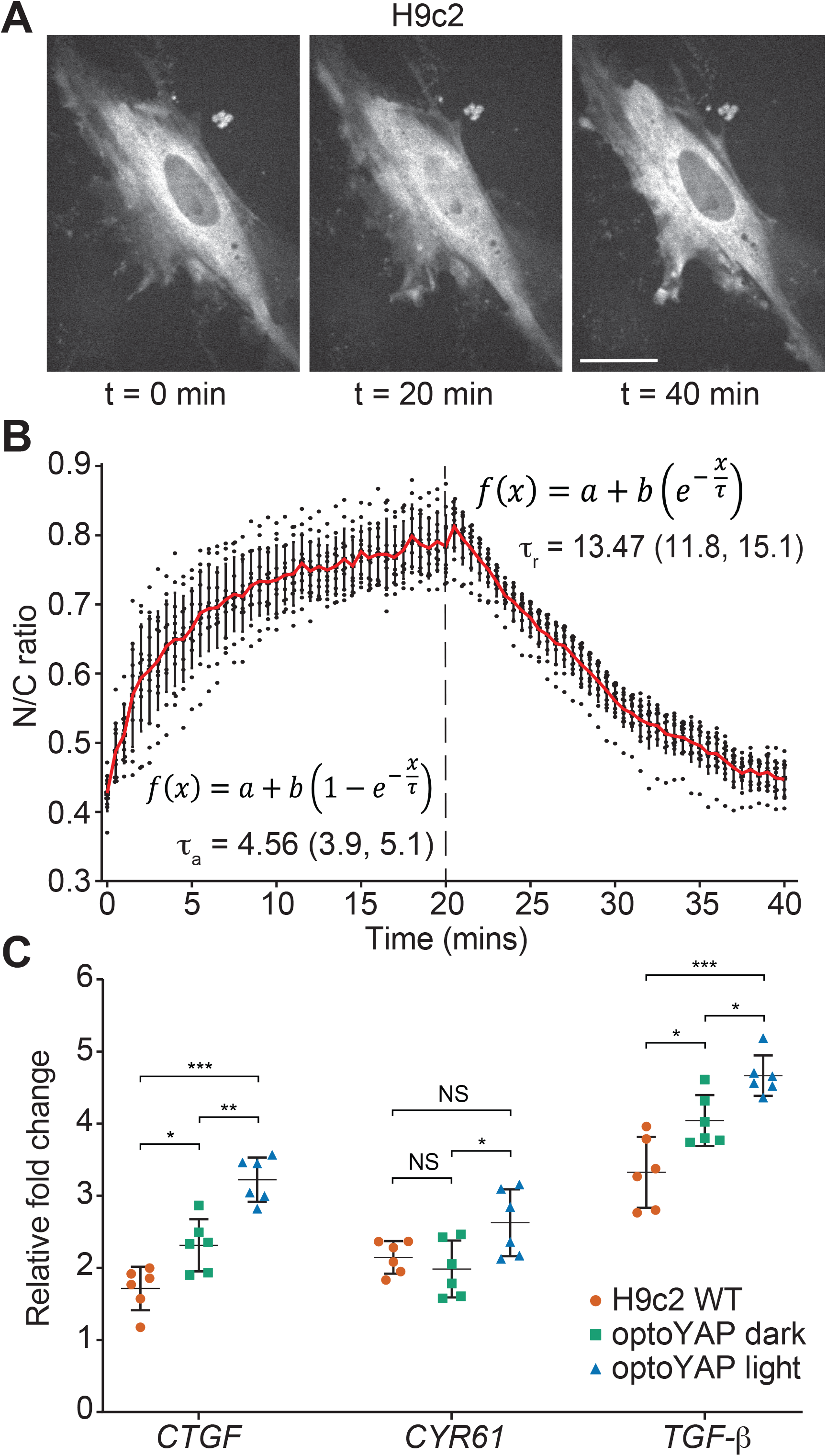
Characterisation of optoYAP in H9c2 cells. **(A)** Representative images of H9c2 cells transfected with optoYAP kept in the dark (t = 0 min), then subjected to pulsatile light activation for 20 min (t = 20 min), followed by recovery in the dark for a further 20 min (t = 40 min). Scale bar, 20 μm. **(B)** Quantification of activation (τ_a_) and recovery (τ_r_) time constant of optoYAP. Red line represents average nuclear/cytoplasmic ratio. Vertical dashed line represents time when 488 nm stimulation ceased. Numbers in brackets represent the 95% confidence interval of τ. n = 12 cells. **(C)** RT-qPCR of YAP downstream genes. Cells were transfected with optoYAP, and transcript levels were assessed after 48 h of activation. Expression levels of *CTGF, CYR61*, and *TGF-β* were normalised to the housekeeping gene *EIF1B* and described as relative fold change. Horizontal bars represent mean and 95% confidence interval from 6 biological replicates from two independent experiments. NS: not significant, \**P* < 0.05, \*\**P* < 10^−2^, \*\*\**P* < 10^−3^.

### Elevated downstream gene expression levels after optoYAP activation

Following the light-dependent nuclear localisation of optoYAP, we examined the functionality of nuclear localised optoYAP by measuring the changes in downstream gene expression levels. We quantified the transcript levels of *CTGF, CYR61*, and *TGF-β* in H9c2 cells after 48 h of pulsatile light activation (Fig. 1C). *CTGF* and *CYR61* are canonical YAP target genes that provide a readout for optoYAP activity in the nucleus (18,29). Additionally, we looked at *TGF-β* transcript levels, an inflammation-related growth factor that is frequently involved in the migration and invasion of tumour cells (47). YAP has also been shown to be a mediator of TGF-*β* signalling (48,49) and the pathway has been implicated in cardiomyocyte proliferation and migration (50,51).

We observed that the expression levels of all three genes probed were significantly upregulated in cells containing activated optoYAP as compared to wildtype (WT) H9c2 (Fig. 1C). However, the transcript levels of *CTGF* and *TGF-β* in transfected cells kept in the dark were also significantly increased in comparison to wildtype controls, though clearly less than when light activated. This suggests that there might be small amounts of nuclear optoYAP even without light activation. However, the upregulation of gene expression across the three genes is significant when comparing cells kept in the light versus dark. This shows that the optoYAP construct is functional when nuclear localised and can trigger expression of downstream target genes.

### optoYAP promotes collective cell migration

We next investigated the role of optoYAP during wound healing, comparing endogenous and optogenetic YAP. Serum-starved cells were seeded in wells 500 μm apart in serum-free media and the “wound” size was quantified every 30 min until complete closure. We observed that optoYAP transfected H9c2 cells kept under pulsatile light activation migrated faster over the wound as compared to transfected cells kept in the dark or untransfected H9c2 (Fig. 2A). Wildtype H9c2 cells took an average of 66 h for complete wound closure, in comparison to 60 h taken for transfected cells in the dark (Fig. 2B). Light activated optoYAP accelerated wound healing, taking an average of 53 h, a 20% improvement on the wildtype cells. Though optoYAP transfected H9c2 cells kept in the dark took a shorter time as compared to wildtype cells, light activation of the optogenetic construct clearly bolstered this behaviour. Extracting the curve slopes in Fig. 2B, we see that the activated optoYAP cells have an increased rate of closure (Fig. 2B inset).

**Figure 2.**
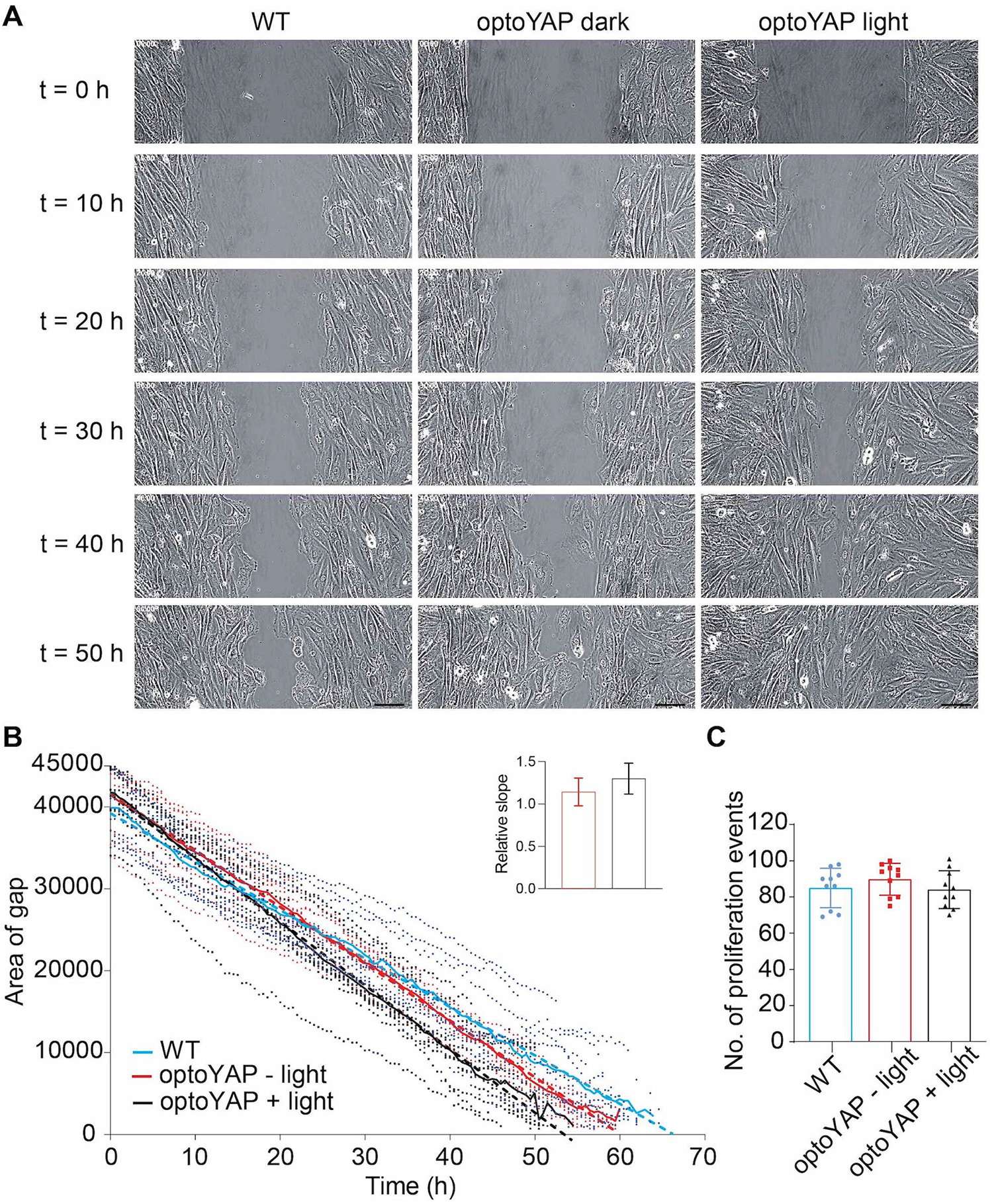
Wound healing assay in H9c2 cells. **(A)** Timelapse images of wildtype H9c2 cells or cells transfected with optoYAP under different light conditions, taken every 30 min until gap completely closes. Representative images shown at 10 h intervals. Scale bars, 100 μm. **(B)** Time taken for H9c2 cells for complete wound healing. Quantification of data in (A) over time. Bold lines represent mean and dashed lines represent linear regression fitted to the curve. n = 10 from two independent experiments. Inset shows the measured slopes under different conditions relative to wildtype (colours as in main plot), error bars are s.d.. **(C)** Number of proliferation events counted during each wound healing experiment in (B). Bars represent mean and error bars represents s.d..

We observed some cell proliferation events during the timelapse, despite the serum starvation before and during the experiment, seen as bright, rounded cell doublets in Fig. 2A. We counted the number of proliferation events that occurred during wound healing and observed that all three conditions had similar number of proliferation events (Fig. 2C). This shows that the differences in the time taken for wound healing observed in Fig. 2B is largely due to cell migration and not proliferation.

We performed the same wound healing experiment in another cell line, MKN28, to exploit the benefit of a CRISPR *YAP*^*-/-*^ background previously reported (52). This cell line enabled us to precisely delineate the effect of optoYAP without any endogenous YAP present that might confound the above results. Previous studies in the same cell line observed a difference between wildtype and *YAP*^*-/-*^ cells in a trans-well migration assay but not during wound healing assays (52). Therefore, we wanted to determine if our optoYAP construct changes the rate of wound closure in these MKN28 *YAP*^*-/-*^ cells.

As expected, the *YAP*^*-/-*^ cells took much longer than the wildtype or optoYAP transfected cells for wound closure, taking 38 h as compared to 33 h for wildtype MKN28 cells (Fig. 3A). This shows that endogenous YAP is required for cell motility during wound closure. In MKN28 *YAP*^*-/-*^ cells transfected with optoYAP, we noticed that those kept in the dark took a faster time for wound closure (30.5 h) in comparison to wildtype cells. Light activated optoYAP transfected *YAP*^*-/-*^ cells had an accelerated rate of wound closure, taking only 29 h for the wound to close completely. This is supported by analysis of the slope of the curves (Fig. 3A inset). These results support the conclusion that endogenous YAP is involved in cell migration and optoYAP further accelerates wound healing in MKN28 cells. Globally, it was noteworthy that the MKN28 cells took a shorter time for wound closure (<40 h) as compared to the H9c2 cells (>50 h).

**Figure 3.**
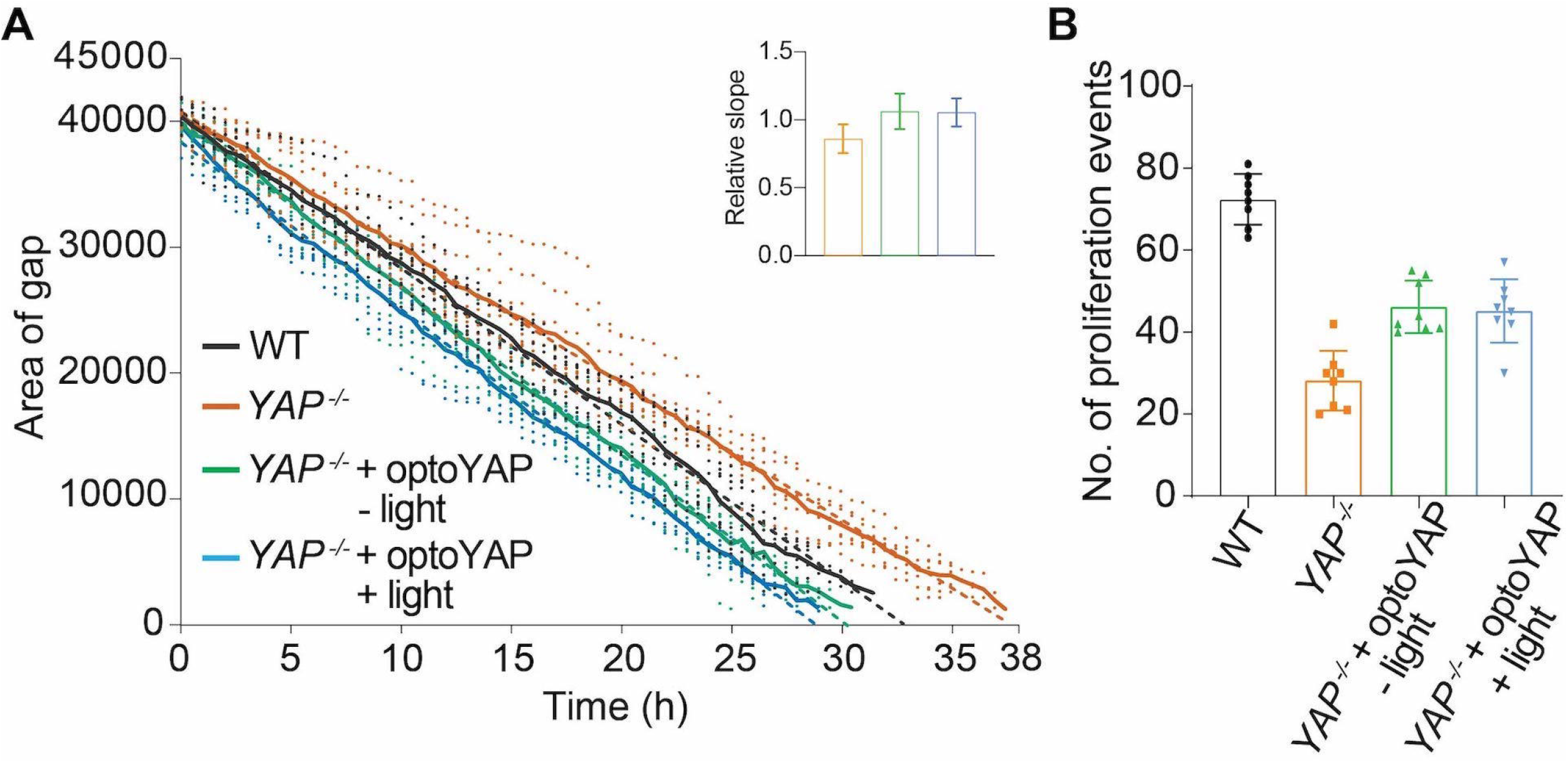
Wound healing assay in MKN28 variants. **(A)** Wildtype (WT) MKN28 and MKN28 *YAP*^*-/-*^ cells transfected with optoYAP were seeded in culture wells 500 μm apart. The size of the gap was tracked every 30 min until it was completely closed. Bold lines represent mean and dashed line represents linear regression fitted to curve. n = 8 from two independent experiments. Inset shows slope of lines relative to wildtype (colours as in main plot), with error bars representing s.d.. **(B)** Number of proliferation events in each experiment in (A). Bars represent mean and error bars represents s.d..

As with the H9c2 cells, we observed several proliferating MKN28 cells in the cell islands during wound closure. We counted the number of proliferation events and compared it amongst the MKN28 variants (Fig. 3B). Unsurprisingly, the *YAP*^*-/-*^ variants had the least amount of proliferation events (28 ± 7), less than half of the wildtype cells (72 ± 6) over the entire time course. Since YAP plays an important role in proliferation, knocking out YAP naturally reduces cell proliferation, which might explain the longer time taken for wound closure in *YAP*^*-/-*^ cells as compared to wildtype. Both optoYAP transfected *YAP*^*-/-*^ cells under light (45 ± 7) or dark (46 ± 6) conditions had similar number of proliferation events, but still less than wildtype cells. Despite less proliferation observed in optoYAP transfected *YAP*^*-/-*^ cells, the time taken for wound closure is shorter than wildtype cells. This shows that optoYAP speeds up the rate of wound closure in *YAP*^*-/-*^ cells, seemingly by inducing cell migration. However, cell proliferation plays a crucial part in wound closure, and it is important to understand the fine interplay between proliferation and migration as revealed here by using CRISPR/Cas9-generated *YAP*^*-/-*^ MKN28 cells.

### Changes in nuclear localisation of optoYAP mutants

Above, we have shown that YAP plays a role in wound healing, including in cardiomyocytes. Phosphorylation of YAP on two ERK activating sites appears to be critical in this response (25), but the dynamics remain poorly understood. To explore this further, we synthesised three phosphomimetic optoYAP mutants by substituting serine to aspartic acid on either S251 or S333, or both. We did this in HEK293T cells due to their ease of transfection over H9c2 or MKN28 cells. We did not observe distinct nuclear localisation differences amongst the three mutants (Fig. 4A). Comparing the basal nuclear localisation of the three mutants, we observed that the double mutant had a significantly higher initial nuclear localisation compared to the S251D mutant, but not the S333D mutant (Fig. 4B). The mutants all displayed clear response to light activation (Fig. 4B-D). They also displayed higher initial nuclear localisation over wildtype optoYAP (Fig. 4E).

**Figure 4.**
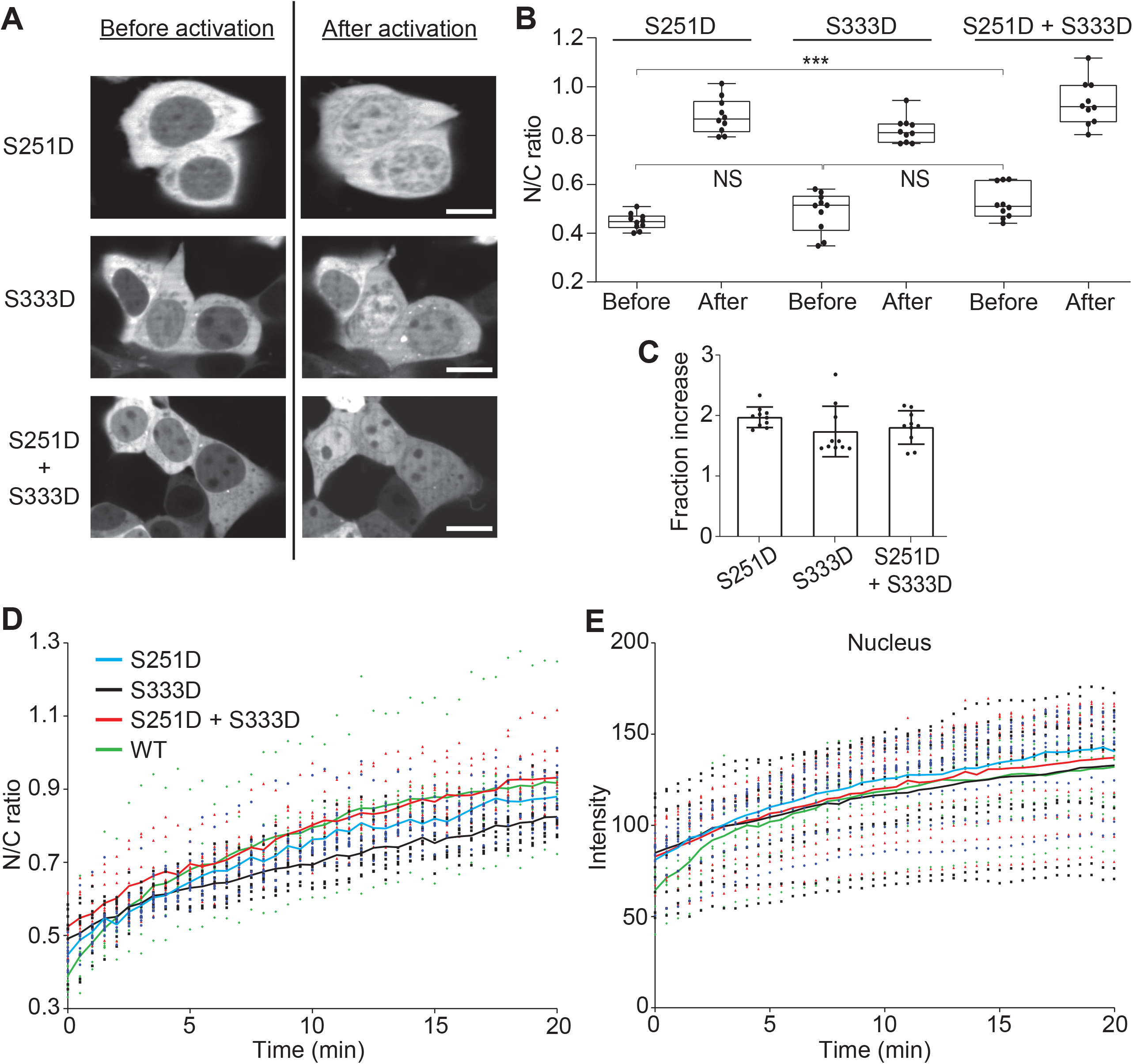
Point mutations of optoYAP at ERK phosphorylation serine residues. **(A)** Three mutants containing phosphomimetic of ERK target residues at S251, S333 or both S251 and S333. Representative images of HEK293T cells transfected with different mutant optoYAP before and after exposure to pulsatile light activation. Scale bars, 10 μm. **(B)** Quantification of cells from (A), measuring nuclear to cytoplasmic ratio of optoYAP before and after light activation for each mutant. NS: not significant, \*\*\**P* < 10^−3^. **(C)** Fraction increase of N/C ratio after light activation. Box plots represent median and 25^th^ to 75^th^ percentiles. Whiskers show minimum and maximum points. **(D)** Relative N/C ratio over time during 488 nm activation amongst optoYAP mutants. **(E)** Absolute intensity values of mCherry signal in nuclei over time. Solid lines represent mean, n = 10 cells taken from three independent experiments.

We then looked at canoncial YAP regulatory sites, such as serine 127 and the C-terminal PDZ-binding motif. We observed that the S127A mutant (referred to as optoYAP S127A) is nuclear localised even before light activation, contrasting significantly with the wildtype cells (Fig. 5A-B). After light activation, we observed that optoYAP S127A accumulated further within the nucleus, doubling the nuclear to cytoplasmic ratio of optoYAP S127A as compared to the dark state (Fig. 5D). This shows that the optoYAP S127A mutant can exert its predominantly nuclear state but optogenetic activation can intensify this ability and increase optoYAP levels in the nucleus.

**Figure 5.**
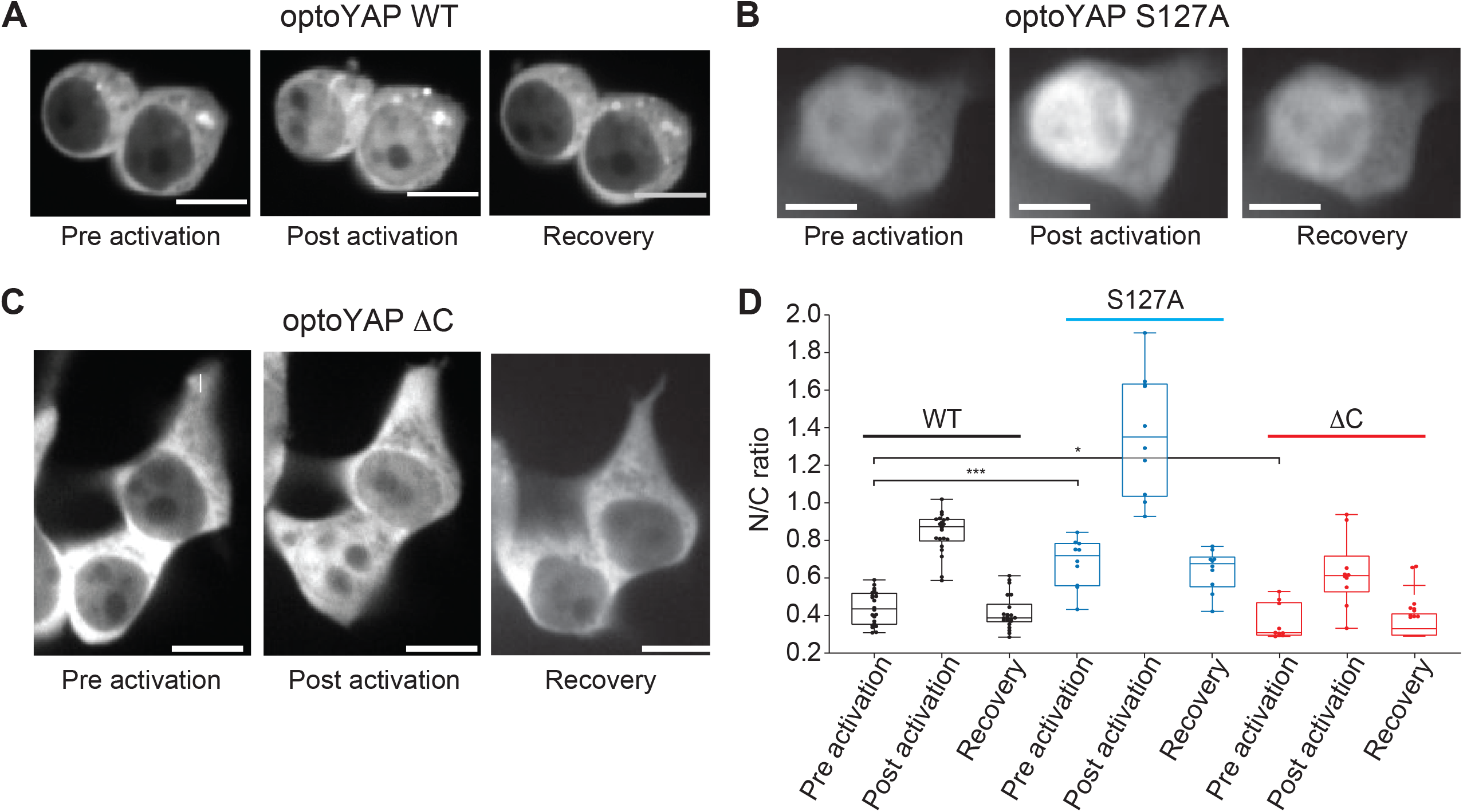
optoYAP variants under light activation protocol. Representative images of mCherry signal in HEK293T cells containing **(A)** wildtype (WT) optoYAP, **(B)** optoYAP with a point mutation at S127A, or **(C)** truncated optoYAP without the C-terminal PDZ-binding motif (optoYAP ΔC). The ‘Pre activation’ panels show cells before exposure to 488 nm light, and after 20 min of pulsatile activation with 488 nm light, the same cells were imaged as ‘Post activation’. Cells were left in the dark for a 20 min and imaged again, under the ‘Recovery’ panel. **(D)** Quantification of nuclear to cytoplasmic ratio of optoYAP in (A-C). Box plots represent median and 25^th^ to 75^th^ percentiles. Whiskers show minimum and maximum points. Scale bars, 10 μm. n = 22 cells for (A), n = 10 cells for (B) and (C), taken from three independent experiments. \**P* < 0.05, \*\*\**P* < 10^−3^.

Deletion of the PDZ-binding motif on YAP causes complete YAP exclusion from the nucleus (32). We synthesised a truncated optoYAP without the PDZ-binding motif, which we refer to as optoYAP ΔC. We observed nuclear exclusion of optoYAP ΔC before exposure to light activation (Fig. 5C). Pulsatile light activation can induce nuclear localisation of optoYAP ΔC, nearly doubling the amount of nuclear localised optoYAP ΔC. The optoYAP ΔC construct returns to the cytoplasm after 20 min of recovery in the dark, similar to the wildtype and S127A mutant. Therefore, activation of the optogenetic tool can overcome mutations that drive YAP out of the nucleus.

## Discussion

Here, we report that optoYAP can accelerate wound healing by promoting cell proliferation and migration in different cell culture systems. We show that activation of YAP can increase the rate of wound repair by 20% in cardiomyocytes. optoYAP, in the absence of endogenous YAP in MKN28 cells, promotes wound healing, accelerating the time taken for closure (Fig. 3A). However, we observed smaller differences in the H9c2 cells between the wildtype and the optoYAP transfected cells (Fig. 2A-B). We further explored the mechanisms that determine the YAP nuclear-cytoplasmic localisation. We demonstrate that the defects in YAP nuclear localisation in the S127A mutant and truncated C-terminal PDZ-binding motif can be compensated through the use of optoYAP activation. Taken together, our results provide strong evidence that: (i) YAP can play an active role in repair beyond simply proliferation; and (ii) optoYAP can regulate YAP activity, even in the presence of different perturbations to YAP, which may be important for wound repair in a range of cell types.

By counting the number of cell proliferation events during wound closure, we observed a few differences between the MKN28 and H9c2 cell lines. We saw that optoYAP increases the amount of cell proliferation in serum starved MKN28 *YAP*^*-/-*^ cells, but not to the wildtype levels (Fig. 3B). However, in MKN28 *YAP*^*-/-*^ cells, the lack of endogenous YAP reduces the amount of proliferation in these cells. Despite transfection with optoYAP, the levels of YAP present in MKN28 *YAP*^*-/-*^ cells might not be the same as endogenous YAP in wildtype cells. Therefore, this might explain the differences between the amount of proliferation in the MKN28 variants. On the contrary in H9c2 cells, activated optoYAP does not alter the amount of proliferation in transfected cells (Fig. 2C). This demonstrates that serum starvation effectively reduces cell proliferation globally. Overall, both cell lines with transfected optoYAP kept in the dark had comparable numbers of proliferation to their light activated counterparts. This shows that despite having similar levels of cell proliferation, light activated optoYAP speeds up the time required for wound closure in both cell lines by enhancing cell migration.

Multiple signalling pathways regulate YAP independently of Hippo, including the EGFR signalling pathway (53). Downstream of EGFR lie the Ras/Raf/MEK/ERK transduction axis, where ERK is the final kinase activator that regulates transcription factors. ERK is a key regulator of regeneration and interplay between ERK and YAP has been shown to be drive YAP activation. By synthesising phosphomimetics of the ERK phosphorylation sites, we observed some nuclear localisation of optoYAP without light activation. Nuclear localisation of YAP phosphorylated at both serine residues have been observed, though a single phosphorylation mark at either S251 or S333 is sufficient to induce nuclear translocation (54,55). However, the double mutant is significantly more nuclear localised than the single mutants (Fig. 4B), which shows the additive effect of phosphorylating both serine residues. Despite the differences observed amongst the three optoYAP mutants, it appears that mutating optoYAP at the two ERBB2 activating phosphorylation sites does not significantly increase nuclear localisation of the optoYAP construct as compared to the wildtype variant.

Additionally, we looked at canonical YAP regulatory sites, S127 and the PDZ-binding motif, that changes the cellular distribution of YAP. Despite all three mutants having an increase in nuclear localisation after light activation, it is evident that the basal level of nuclear localisation is not equal. The optoYAP S127A mutant (0.68 ± 0.13) has a significantly higher initial nuclear concentration than wildtype (0.44 ± 0.09), since the S127A mutation causes optoYAP to be predominantly nuclear. Conversely, the optoYAP ΔC mutant (0.36 ± 0.09) has a significantly lower nuclear localisation compared to wildtype (0.44 ± 0.09), which is expected as deletion of the PDZ-binding motif prevents optoYAP from translocating into the nucleus (32). Although the basal levels differ for the three variants, nuclear localisation doubles for all optoYAP variants and decreases back to their respective basal levels after recovery in the dark (Fig. 5D). This shows that the optogenetic domains can promote further nuclear translocation despite mutations that already boosts (S127A) or inhibit (ΔC) nuclear localisation.

The rate of nuclear localisation of the two S127A and ΔC optoYAP mutants have not been studied, which could shed light on the dynamics of nuclear shuttling in the optoYAP mutants. The rate of nuclear import or export might differ drastically between the optoYAP variants, depending on the mutation. We hypothesise that the activating S127A mutation increases the rate of nuclear import but does not affect nuclear export rates, while the negative ΔC mutant increases the rate of nuclear export but maintains similar import rates as wildtype. Regardless, it is evident that mutations in the S127 or PDZ-binding motif changes the initial localisation of optoYAP, but the activated optogenetic construct can supersede the basal distribution and induce optoYAP nuclear localisation.

Finally, we note that several optogenetic YAP constructs have been published recently, with different YAP isoforms and activation protocols. Illes *et al*. first published a tool to control the subcellular localisation of YAP by tagging it with an optogenetic NLS, shuttling YAP between the cytoplasm and the nucleus (56). The authors used the human YAP1 gene without specifying the details of the splice isoform but demonstrated that activated YAP increases proliferation in 2D and 3D cultures and induces invasion in cancer spheroids (56). Dowbaj *et al*. presented optogenetic YAP1, using the human YAP1-2γ isoform (57), which contains exon 6 and two WW domains (58), tethering optogenetic YAP and TAZ to the mitochondria. Their tool allows the study of protein dynamics by describing the rate of nuclear entry and exit but does not demonstrate downstream functionality of nuclear YAP (57). In our previous work, we reported a human YAP1-1δ optogenetic tool that translocates between the cytoplasm and nucleus, demonstrating functional outputs *in vitro* and *in vivo* (40). Most recently, Meyer *et al*. have developed a mouse YAP1 tool. They are able to regulate the levels of pluripotency factors Oct4 and Nanog, and hence control the renewal of a naïve stem cell state or preferential differentiation into the mesendodermal state (59). The various YAP isoforms signal differently, where those without exon 6, alpha and beta, can form heterodimers with TAZ (60). Additionally, different isoforms have unique and common functions and activate distinct downstream transcriptional programmes (61). Taken together, the isoforms of YAP matter when describing optogenetic constructs, given that each isoform can have unique downstream effects.

The role of YAP in wound healing has been investigated previously, where nuclear localisation of YAP is observed during healing (62,63). We demonstrate that optoYAP can accelerate wound healing as compared to endogenous YAP, providing a tight spatiotemporal control over its activity. It has been shown that H9c2 cells respond to substrate stiffness and topography, promoting cell differentiation on a modified extracellular matrix (ECM) synthesized from fibroblast-derived matrix (64). With our optogenetic tool, it will be interesting to investigate cell migration in an environment that mimics physiological conditions by seeding cells on permissive ECM, or a stiffened matrix that imitates cholesterol plaques, and tracking wound healing under such conditions, which will benefit future studies on regeneration.

## Abbreviations

cDNA: complementary DNA
Ct: threshold cycle
DMEM: Dulbecco’s modified Eagle’s medium
ECM: extracellular matrix
EGFR: epidermal growth factor receptor
FBS: fetal bovine serum
NA: numerical aperture
NEAA: non-essential amino acids
NFW: nuclease-free water
NLS: nuclear localisation sequence
ROI: region of interest
RT-qPCR: quantitative real-time polymerase chain reaction
SDM: site-directed mutagenesis
STR: short tandem repeat
WT: wildtype

## Declarations

### Ethics approval and consent to participate

Not applicable

### Consent for publication

Not applicable

### Availability of data and materials

The datasets used and/or analysed during this study are available from the corresponding author on reasonable request.

### Authors’ contributions

PJYT, MS and TES conceived the study and design of the work. PJYT performed the experiments. PJYT and TES did the data analysis. PJYT and TES drafted the work with all authors approving the final manuscript.

## Acknowledgements

We thank Michael Sheetz, in whose lab the work on developing the optogenetic YAP construct was initiated, and Manvendra Singh for comments on the manuscript.

## References

1. Murry CE, Reinecke H, Pabon LM. Regeneration gaps: observations on stem cells and cardiac repair. J Am Coll Cardiol. 2006 May;47(9):1777–85.

2. Sedmera D, Reckova M, DeAlmeida A, Coppen SR, Kubalak SW, Gourdie RG, et al. Spatiotemporal pattern of commitment to slowed proliferation in the embryonic mouse heart indicates progressive differentiation of the cardiac conduction system. Anat Rec A Discov Mol Cell Evol Biol. 2003 Sep;274(1):773–7.

3. Steinhauser ML, Lee RT. Regeneration of the heart. EMBO Mol Med. 2011 Dec 1;3(12):701–12.

4. Porrello ER, Mahmoud AI, Simpson E, Hill JA, Richardson JA, Olson EN, et al. Transient Regenerative Potential of the Neonatal Mouse Heart. Science (1979). 2011;331(6020):1078–80.

5. Poss KD, Wilson LG, Keating MT. Heart Regeneration in Zebrafish. Science (1979). 2002 Dec 13;298(5601):2188–90.

6. Chong JJH, Yang X, Don CW, Minami E, Liu YW, Weyers JJ, et al. Human embryonic-stem-cell-derived cardiomyocytes regenerate non-human primate hearts. Nature. 2014 Jun;510(7504):273–7.

7. Liu YW, Chen B, Yang X, Fugate JA, Kalucki FA, Futakuchi-Tsuchida A, et al. Human embryonic stem cell-derived cardiomyocytes restore function in infarcted hearts of non-human primates. Nat Biotechnol. 2018 Aug;36(7):597–605.

8. Xin M, Kim Y, Sutherland LB, Murakami M, Qi X, McAnally J, et al. Hippo pathway effector Yap promotes cardiac regeneration. Proceedings of the National Academy of Sciences. 2013 Aug 20;110(34):13839–44.

9. Heallen T, Morikawa Y, Leach J, Tao G, Willerson JT, Johnson RL, et al. Hippo signaling impedes adult heart regeneration. Development. 2013 Dec;140(23):4683–90.

10. Leach JP, Heallen T, Zhang M, Rahmani M, Morikawa Y, Hill MC, et al. Hippo pathway deficiency reverses systolic heart failure after infarction. Nature. 2017 Oct;550(7675):260–4.

11. Panciera T, Azzolin L, Cordenonsi M, Piccolo S. Mechanobiology of YAP and TAZ in physiology and disease. Nat Rev Mol Cell Biol. 2017;18(12):758–70.

12. Fu V, Plouffe SW, Guan KL. The Hippo pathway in organ development, homeostasis, and regeneration. Curr Opin Cell Biol. 2017;49:99–107.

13. Basu S, Totty NF, Irwin MS, Sudol M, Downward J. Akt phosphorylates the Yes-Associated Protein, YAP, to induce interaction with 14-3-3 and attenuation of p73-mediated apoptosis. Mol Cell. 2003;11(1):11–23.

14. Kanai F, Marignani PA, Sarbassova D, Yagi R, Hall RA, Donowitz M, et al. TAZ: a novel transcriptional co-activator regulated by interactions with 14-3-3 and PDZ domain proteins. EMBO J. 2000 Dec 15;19(24):6778–91.

15. Zhao B, Li L, Tumaneng K, Wang CY, Guan KL. A coordinated phosphorylation by Lats and CK1 regulates YAP stability through SCFβ-TRCP. Genes Dev. 2010;24(1):72–85.

16. Vassilev A, Kaneko KJ, Shu H, Zhao Y, DePamphilis ML. TEAD/TEF transcription factors utilize the activation domain of YAP65, a Src/Yes-associated protein localized in the cytoplasm. Genes Dev. 2001 May 15;15(10):1229–41.

17. Zhang H, Liu CY, Zha ZY, Zhao B, Yao J, Zhao S, et al. TEAD Transcription Factors Mediate the Function of TAZ in Cell Growth and Epithelial-Mesenchymal Transition. Journal of Biological Chemistry [Internet]. 2009 May 15;284(20):13355–62. Available from: https://doi.org/10.1074/jbc.M900843200

18. Zhao B, Ye X, Yu J, Li L, Li W, Li S, et al. TEAD mediates YAP-dependent gene induction and growth control. Genes Dev. 2008;22(14):1962–71.

19. Yu FX, Zhao B, Guan KL. Hippo Pathway in Organ Size Control, Tissue Homeostasis, and Cancer. Cell. 2015;163(4):811–28.

20. Moya IM, Halder G. Hippo–YAP/TAZ signalling in organ regeneration and regenerative medicine. Nat Rev Mol Cell Biol. 2019;20:211–26.

21. Lai JKH, Toh PJY, Cognart HA, Chouhan G, Saunders TE. DNA-damage induced cell death in yap1;wwtr1 mutant epidermal basal cells. Elife. 2022 May 1;11.

22. Wen X, Jiao L, Tan H. MAPK/ERK Pathway as a Central Regulator in Vertebrate Organ Regeneration. Int J Mol Sci. 2022;23(3).

23. De Simone A, Evanitsky MN, Hayden L, Cox BD, Wang J, Tornini VA, et al. Control of osteoblast regeneration by a train of Erk activity waves. Nature. 2021 Feb;590(7844):129–33.

24. Bassat E, Mutlak YE, Genzelinakh A, Shadrin IY, Baruch Umansky K, Yifa O, et al. The extracellular matrix protein agrin promotes heart regeneration in mice. Nature. 2017;547(7662):179–84.

25. Aharonov A, Shakked A, Umansky KB, Savidor A, Genzelinakh A, Kain D, et al. ERBB2 drives YAP activation and EMT-like processes during cardiac regeneration. Nat Cell Biol. 2020;22(11):1346–56.

26. Oka T, Mazack V, Sudol M. Mst2 and Lats kinases regulate apoptotic function of Yes kinase-associated protein (YAP). Journal of Biological Chemistry. 2008;283(41):27534–46.

27. Ege N, Dowbaj AM, Jiang M, Howell M, Hooper S, Foster C, et al. Quantitative Analysis Reveals that Actin and Src-Family Kinases Regulate Nuclear YAP1 and Its Export. Cell Syst. 2018/06/18. 2018 Jun 27;6(6):692-708.e13.

28. Zhao B, Wei X, Li W, Udan RS, Yang Q, Kim J, et al. Inactivation of YAP oncoprotein by the Hippo pathway is involved in cell contact inhibition and tissue growth control. Genes Dev. 2007;21:2747–61.

29. Zhang H, Pasolli HA, Fuchs E. Yes-associated protein (YAP) transcriptional coactivator functions in balancing growth and differentiation in skin. Proceedings of the National Academy of Sciences. 2011;108(6):2270–5.

30. Lamar JM, Stern P, Liu H, Schindler JW, Jiang ZG, Hynes RO. The Hippo pathway target, YAP, promotes metastasis through its TEAD-interaction domain. Proc Natl Acad Sci U S A. 2012 Sep;109(37):E2441–50.

31. Oka T, Remue E, Meerschaert K, Vanloo B, Boucherie C, Gfeller D, et al. Functional complexes between YAP2 and ZO-2 are PDZ domain-dependent, and regulate YAP2 nuclear localization and signalling. Biochemical Journal. 2010;432(3):461–78.

32. Oka T, Sudol M. Nuclear localization and pro-apoptotic signaling of YAP2 require intact PDZ-binding motif. Genes to Cells. 2009;14(5):607–15.

33. Mason DE, Collins JM, Dawahare JH, Nguyen TD, Lin Y, Voytik-Harbin SL, et al. YAP and TAZ limit cytoskeletal and focal adhesion maturation to enable persistent cell motility. Journal of Cell Biology. 2019 Feb 8;218(4):1369–89.

34. D’Angelo E, Lindoso RS, Sensi F, Pucciarelli S, Bussolati B, Agostini M, et al. Intrinsic and Extrinsic Modulators of the Epithelial to Mesenchymal Transition: Driving the Fate of Tumor Microenvironment. Vol. 10, Frontiers in Oncology. Frontiers Media S.A.; 2020.

35. Morikawa Y, Zhang M, Heallen T, Leach J, Tao G, Xiao Y, et al. Actin cytoskeletal remodeling with protrusion formation is essential for heart regeneration in Hippo-deficient mice. Sci Signal. 2015 May;8(375):ra41.

36. Mia MM, Cibi DM, Ghani Saba, Singh A, Tee N, Sivakumar V, et al. Loss of Yap/Taz in cardiac fibroblasts attenuates adverse remodelling and improves cardiac function. Cardiovasc Res [Internet]. 2022 May 15;118(7):1785–804. Available from: https://doi.org/10.1093/cvr/cvab205

37. Zhou J. An emerging role for Hippo-YAP signaling in cardiovascular development. J Biomed Res. 2014 Jul;28(4):251–4.

38. Fenno L, Yizhar O, Deisseroth K. The development and application of optogenetics. Annu Rev Neurosci [Internet]. 2011;34:389–412. Available from: https://pubmed.ncbi.nlm.nih.gov/21692661

39. Krueger D, Izquierdo E, Viswanathan R, Hartmann J, Pallares Cartes C, de Renzis S. Principles and applications of optogenetics in developmental biology. Development [Internet]. 2019 Oct 22;146(20):dev175067. Available from: https://doi.org/10.1242/dev.175067

40. Toh PJY, Lai JKH, Hermann A, Destaing O, Sheetz MP, Sudol M, et al. Optogenetic control of YAP cellular localisation and function. EMBO Rep [Internet]. 2022 Sep 5;23(9):e54401. Available from: https://doi.org/10.15252/embr.202154401

41. Kuznetsov A v, Javadov S, Sickinger S, Frotschnig S, Grimm M. H9c2 and HL-1 cells demonstrate distinct features of energy metabolism, mitochondrial function and sensitivity to hypoxia-reoxygenation. Biochimica et Biophysica Acta (BBA) - Molecular Cell Research [Internet]. 2015;1853(2):276–84. Available from: https://www.sciencedirect.com/science/article/pii/S0167488914004042

42. Watkins SJ, Borthwick GM, Arthur HM. The H9C2 cell line and primary neonatal cardiomyocyte cells show similar hypertrophic responses in vitro. In Vitro Cell Dev Biol Anim. 2011 Feb;47(2):125–31.

43. Li C, Huang Z, Wang RK. Elastic properties of soft tissue-mimicking phantoms assessed by combined use of laser ultrasonics and low coherence interferometry. Opt Express. 2011 May;19(11):10153–63.

44. Nitta T, Haga H, Kawabata K, Abe K, Sambongi T. Comparing microscopic with macroscopic elastic properties of polymer gel. Ultramicroscopy. 2000;82(1):223–6.

45. Blainey P, Krzywinski M, Altman N. Replication. Nat Methods. 2014;11(9):879–80.

46. Ho J, Tumkaya T, Aryal S, Choi H, Claridge-Chang A. Moving beyond P values: data analysis with estimation graphics. Nat Methods. 2019;16(7):565–6.

47. Liu Z, Yi L, Du M, Gong G, Zhu Y. Overexpression of TGF-β enhances the migration and invasive ability of ectopic endometrial cells via ERK/MAPK signaling pathway. Exp Ther Med. 2019;17(6):4457–64.

48. Szeto SG, Narimatsu M, Lu M, He X, Sidiqi AM, Tolosa MF, et al. YAP/TAZ are mechanoregulators of TGF-b-smad signaling and renal fibrogenesis. Journal of the American Society of Nephrology. 2016;27(10):3117–28.

49. Qin Z, Xia W, Fisher GJ, Voorhees JJ, Quan T. YAP/TAZ regulates TGF-β/Smad3 signaling by induction of Smad7 via AP-1 in human skin dermal fibroblasts. Cell Communication and Signaling. 2018;16(1):18.

50. Sorensen DW, van Berlo JH. The Role of TGF-β Signaling in Cardiomyocyte Proliferation. Curr Heart Fail Rep. 2020 Oct;17(5):225–33.

51. Peng Y, Wang W, Fang Y, Hu H, Chang N, Pang M, et al. Inhibition of TGF-β/Smad3 Signaling Disrupts Cardiomyocyte Cell Cycle Progression and Epithelial– Mesenchymal Transition-Like Response During Ventricle Regeneration. Front Cell Dev Biol. 2021;9.

52. Qiao Y, Chen J, Lim YB, Finch-Edmondson ML, Seshachalam VP, Qin L, et al. YAP regulates actin dynamics through ARHGAP29 and promotes metastasis. Cell Rep [Internet]. 2017;19(8):1495–502. Available from: http://www.sciencedirect.com/science/article/pii/S2211124717306046

53. Reddy BVVG, Irvine KD. Regulation of Hippo Signaling by EGFR-MAPK Signaling through Ajuba Family Proteins. Dev Cell. 2013;24(5):459–71.

54. Bui DA, Lee W, White AE, Harper JW, Schackmann RCJ, Overholtzer M, et al. Cytokinesis involves a nontranscriptional function of the Hippo pathway effector YAP. Sci Signal. 2016 Mar 1;9(417):ra23–ra23.

55. Yang S, Zhang L, Liu M, Chong R, Ding SJ, Chen Y, et al. CDK1 Phosphorylation of YAP Promotes Mitotic Defects and Cell Motility and Is Essential for Neoplastic Transformation. Cancer Res. 2013 Nov 14;73(22):6722–33.

56. Illes B, Fuchs A, Gegenfurtner F, Ploetz E, Zahler S, Vollmar AM, et al. Spatio-selective activation of nuclear translocation of YAP with light directs invasion of cancer cell spheroids. iScience [Internet]. 2021;24(3):102185. Available from: https://www.sciencedirect.com/science/article/pii/S258900422100153X

57. Dowbaj AM, Jenkins RP, Williamson D, Heddleston JM, Ciccarelli A, Fallesen T, et al. An optogenetic method for interrogating YAP1 and TAZ nuclear–cytoplasmic shuttling. J Cell Sci [Internet]. 2021 Jul 9;134(13):jcs253484. Available from: https://doi.org/10.1242/jcs.253484

58. Gaffney CJ, Oka T, Mazack V, Hilman D, Gat U, Muramatsu T, et al. Identification, basic characterization and evolutionary analysis of differentially spliced mRNA isoforms of human YAP1 gene. Gene. 2012 Nov;509(2):215–22.

59. Meyer K, Lammers NC, Bugaj LJ, Garcia HG, Weiner OD. Decoding of YAP levels and dynamics by pluripotency factors. Available from: https://doi.org/10.1101/2022.10.17.512504

60. Ben C, Wu X, Takahashi-Kanemitsu A, Knight CT, Hayashi T, Hatakeyama M. Alternative splicing reverses the cell-intrinsic and cell-extrinsic pro-oncogenic potentials of YAP1. Journal of Biological Chemistry [Internet]. 2020;295(41):13965–80. Available from: https://www.sciencedirect.com/science/article/pii/S0021925817497961

61. Vrbský J, Vinarský V, Perestrelo AR, de La Cruz JO, Martino F, Pompeiano A, et al. Evidence for discrete modes of YAP1 signaling via mRNA splice isoforms in development and diseases. Genomics [Internet]. 2021;113(3):1349–65. Available from: https://www.sciencedirect.com/science/article/pii/S0888754321001014

62. Elbediwy A, Vincent-Mistiaen ZI, Spencer-Dene B, Stone RK, Boeing S, Wculek SK, et al. Integrin signalling regulates YAP and TAZ to control skin homeostasis. Development. 2016 May 15;143(10):1674–87.

63. Lee MJ, Byun MR, Furutani-Seiki M, Hong JH, Jung HS. YAP and TAZ Regulate Skin Wound Healing. Journal of Investigative Dermatology. 2014;134(2):518–25.

64. Suhaeri M, Subbiah R, Van SY, Du P, Kim IG, Lee K, et al. Cardiomyoblast (h9c2) differentiation on tunable extracellular matrix microenvironment. Tissue Eng Part A. 2015 Jun;21(11–12):1940–51.

